# CauloKO: an ordered transposon mutant library in *Caulobacter crescentus*

**DOI:** 10.1101/2022.11.02.514973

**Authors:** Gabriel M. Moore, Justin G. Ramos, Benjamin P. Bratton, Zemer Gitai

## Abstract

Genetic screens are powerful approaches to unveiling new biological insight and ordered redundant transposon libraries have emerged as a primary tool for performing screens of known genetic saturation. Newer sequencing methods based on combinatorial pooling have lowered the cost and time required to generate these libraries. *Caulobacter crescentus* is a gramnegative bacterium that has served as a model for understanding bacterial physiology with a myriad of genetic tools. To add to this collection of tools, we created CauloKO - the first ordered, transposon library in *C. crescentus*. CauloKO includes insertion mutants in 86% of all non-essential genes and 77% of all open reading frames of strain CB15. CauloKO insertion mutants were validated using Sanger sequencing. We also present phenotypic analysis of the CauloKO library using a crystal violet screen for biofilm mutants, which both confirmed previous results and identified new mutants for future studies. This combined approach revealed that the CauloKO library shows promise for screening applications, particularly for phenotypes that require monoclonal populations of cells.

## Introduction

Our understanding of microbiological processes heavily relies on genetic screens. Transposon screens are particularly powerful because transposons typically result in loss-of-function mutations and mapping their insertion sites is fast and easy. Classical genetic screens introduce random transposon mutagenesis to a population of cells and then examine the resulting colonies for a particular phenotype, or lack thereof [1]. While this method has the advantage of being quick and easy for identifying individual mutations, Poisson statistics make it difficult to achieve saturation and ensure every open reading frame is disrupted [2]. On the other end of the spectrum, high-density TnSeq approaches are a powerful way to study the entirety of the functional genome [3]. However, TnSeq studies are performed in complex communities such that they cannot be applied to clonal populations, and it is often difficult to retrieve specific transposon insertions for further studies. These limitations have been overcome by ordered transposon libraries that attempt to include clonal isolates of as many mutants as possible that can tolerate a transposon insertion [4–8]. Several clinically-relevant bacterial species have had libraries constructed in at least one strain [4–8], but the significant cost and effort typically required to generate these libraries has limited their widespread application.

Recently, strategies such as Knockout Sudoku and Cartesian Pooling-Coordinate Sequencing (CP-CSeq) have been developed for rapid determination of the identity and location of transposon mutants in a library with minimal resources [10–13]. These strategies rely on arraying transposon mutants in 96-well plates into matrices that give each mutant a set of coordinates that can be determined in custom sequencing pipelines. These methods of combinatorial pooling and decoding of transposon mutants has led to an increase in the number of ordered transposon libraries available, including *Mycobacterium bovis, Shewanella oneidensis*, and *Bacteroides thetaiotaomicron* [12–14].

*Caulobacter crescentus* is a gram-negative, alpha-proteobacterium predominantly isolated and associated with aqueous and rhizosphere environments [15–16]. *C. crescentus* has emerged as a powerful model system for a wide range of important behaviors including its canonical “crescent” cell shape, cell cycle, and polarity [17–19]. Recent studies have also suggested that *C. crescentus* can function as an animal pathogen [20]. While many of these processes have been studied using classical genetic techniques, the continued discovery of novel determinants in *C. crescentus* is aided by a number of tools generated in the organism, including a cosmid library [21], a fluorescently-tagged localisome library of open reading frames (ORFs) [22], and a web-based visualizer of all published high-throughput genomic data [23]. At present, however, there is no ordered transposon library to facilitate genetic screens that require monoclonal populations of cells or to provide a resource of mutants-of-interest for genetic analysis.

Given the many resources that already exist in *C. crescentus* and relative ease of library generation with modern sequencing methods, we sought to create and assemble an ordered transposon library in *C. crescentus*. We used a combination of tools from the Knockout Sudoku pipeline and a homebrewed sequencing analysis pipeline to generate CauloKO, a library with 77% coverage of predicted ORFs and 86% of non-essential genes in the *C. crescentus* genome. To validate the usefulness of CauloKO as a tool, we phenotypically characterized this library by using a crystal violet screen. We identified biofilm mutants that were both consistent with past work and highlighted novel candidates for further exploration. Our work thus adds to the wealth of tools available for understanding *Caulobacter* physiology in particular and bacterial biology in general.

## Results and Discussion

### Transposon mutagenesis and construction of initial mutant collection

We chose to use the environmentally-isolated *C. crescentus* CB15 background as the parental strain for our ordered library, as it retains holdfasts and other phenotypes that have been lost in lab-adapted strains [24]. To generate transposon insertions, we mated *C. crescentus* CB15 with pMiniHimar1-possessing *E. coli* WM3064, and selected colonies by plating on antibiotic agar plates with kanamycin and nalidixic acid (Figure 1). Kanamycin-resistance selects for the presence of the miniHimar1 transposon, and *C. crescentus* is naturally-resistant to nalidixic acid thereby eliminating *E. coli* contamination [25].

**Figure 1:**
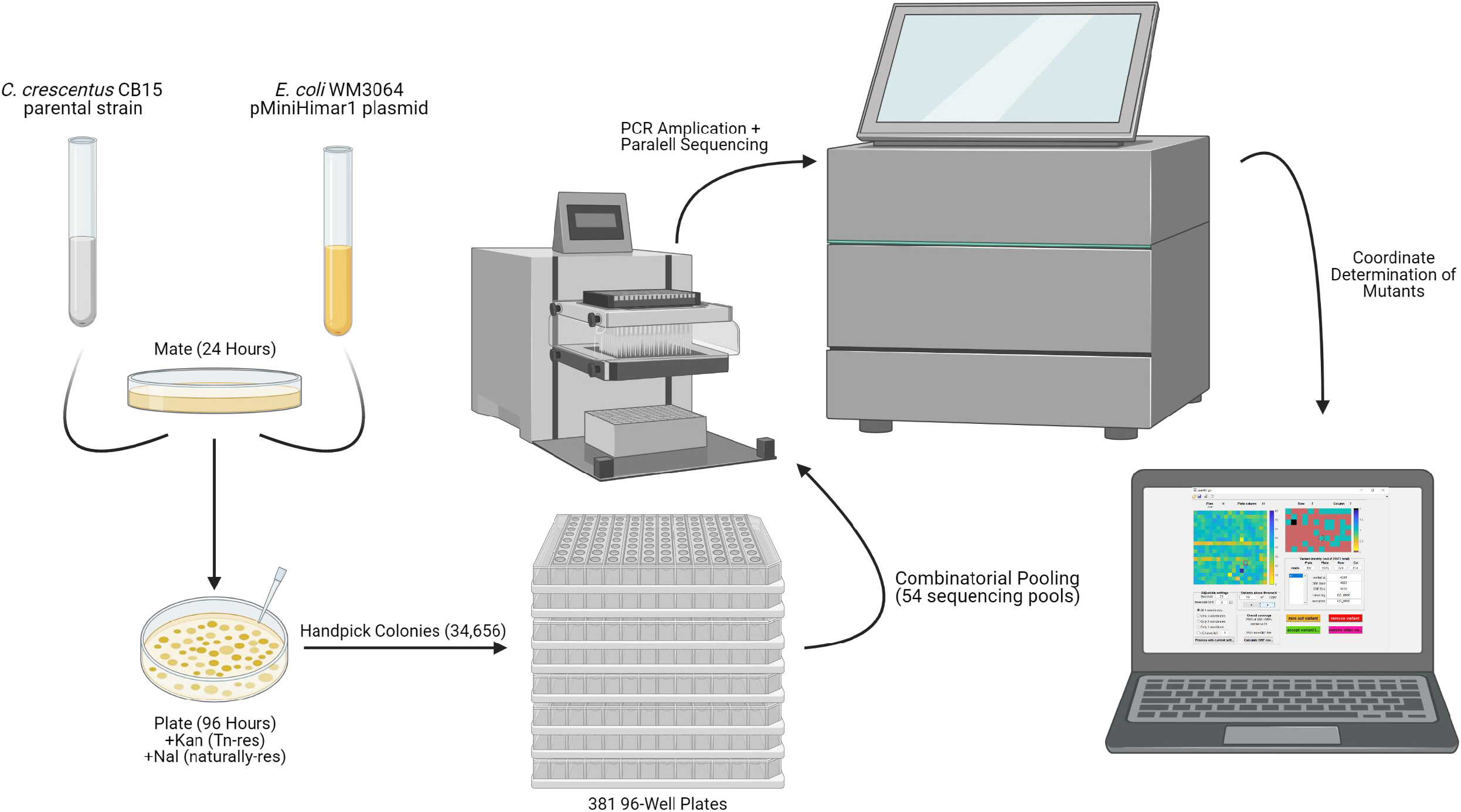
Methodology for physical construction of the *C. crescentus* CB15 CauloKO library. *C. crescentus* CB15 cells were mated with pMiniHimar1 containing E. coli WM3604 and resulting colonies were handpicked in 96-well plates. Plates were pooled into 54 sequencing pots that were barcoded and PCR amplified before sequencing. Determination of the coordinates for sequenced mutants was then determined using the CauloKO GUI generated in this study.

The next step was to pick a large number of *C. crescentus* transposon-containing colonies with the goal of saturating the CB15 genome. CB15 contains 3,767 predicted ORFs. While we have essentiality data for many of these ORFs based on high-density TnSeq analysis of the lab *C. crescentus* strain, NA1000 [26], we calculated an upper estimate for the number of transposon mutants we should include by assuming that all genes have the potential to tolerate transposon mutagenesis. A Monte Carlo-based simulation predicted that 34,656 transposon mutants would cover 99% of all ORFs (Figure 1, Supplementary Figure 1). We note that the Monte Carlo estimate is superior to a Poisson estimate for saturation as it takes into account the length of each gene, so as to not overestimate insertions in long genes nor underestimate insertions in shorter genes [13]. We thus hand-picked 34,656 kanamycin-resistant CB15 colonies from our pMiniHimar1 mating into 361 multi-well plates, each with 96-wells.

We next used the Knockout Sudoku approach to pool our 361 plates into sequencing pools [13]. In brief, the 361 plates were organized into a 19×19 higher-order array. From this array we generated 58 mutant pools, including 12 “column” pools and 8 “row” pools (pooling all of the mutants from each column and row across all 361 plates), as well as 19 “plate column” pools and 19 “plate row” pools (pooling all of the mutants from the entire 96-well plates in each of the 19×19 higher-order plate-columns and plate-rows). These 54 pools enable us to identify each specific mutant from only four coordinates: columns of the plates (1-12), rows of the plates (A-H), and the plate column and plate row within the 19×19 array. Each pool was PCR-amplified, barcoded, and sequenced by Illumina Sequencing.

To ensure no major genomic differences occurred in the time it took to pick, pool, and move all mutants to frozen storage, we performed full genome sequencing of the parental strain and a representative transposon mutant from the collection. A total of 45 SNPs were identified in the parental strain compared to the reference genome for *C. crescentus* CB15 (Supplementary Table 1) [27]. Of these SNPs, 42 were shared in the transposon mutant along with an additional 3 not found in the parental strain. The majority of these SNPs were in intergenic regions or represented silent mutations, and thus unlikely to have an effect on gene products. Thus, the parental strain for library construction reflects a representative CB15 strain and shares nearly identical genomic similarity to the transposon mutants aside from their respective transposon insertion (Supplementary Table 1).

### Sequencing analysis of the initial mutant collection and assembly of the ordered CauloKO library

Sequence analysis of the transposon library pools revealed that our library achieved broad coverage of the *C. crescentus* genome. We successfully identified 22,219 insertions across the CB15 genome, with an insertion rate of ~1 transposon per 278 base pairs (bps). As expected, operons that have previously been shown to be essential were under-represented, as in the case of the essential *nuo* operon for NADH dehydrogenase whose insertion rate was 1 transposon per 1250 bps (Figure 2). Similarly, we observed no insertions in the essential ribosome subunit operon found from *CC1245-CC1273*. Overall, the initial collection of 22,219 insertions included at least one transposon insertion in 85.6% of all predicted genes.

**Figure 2:**
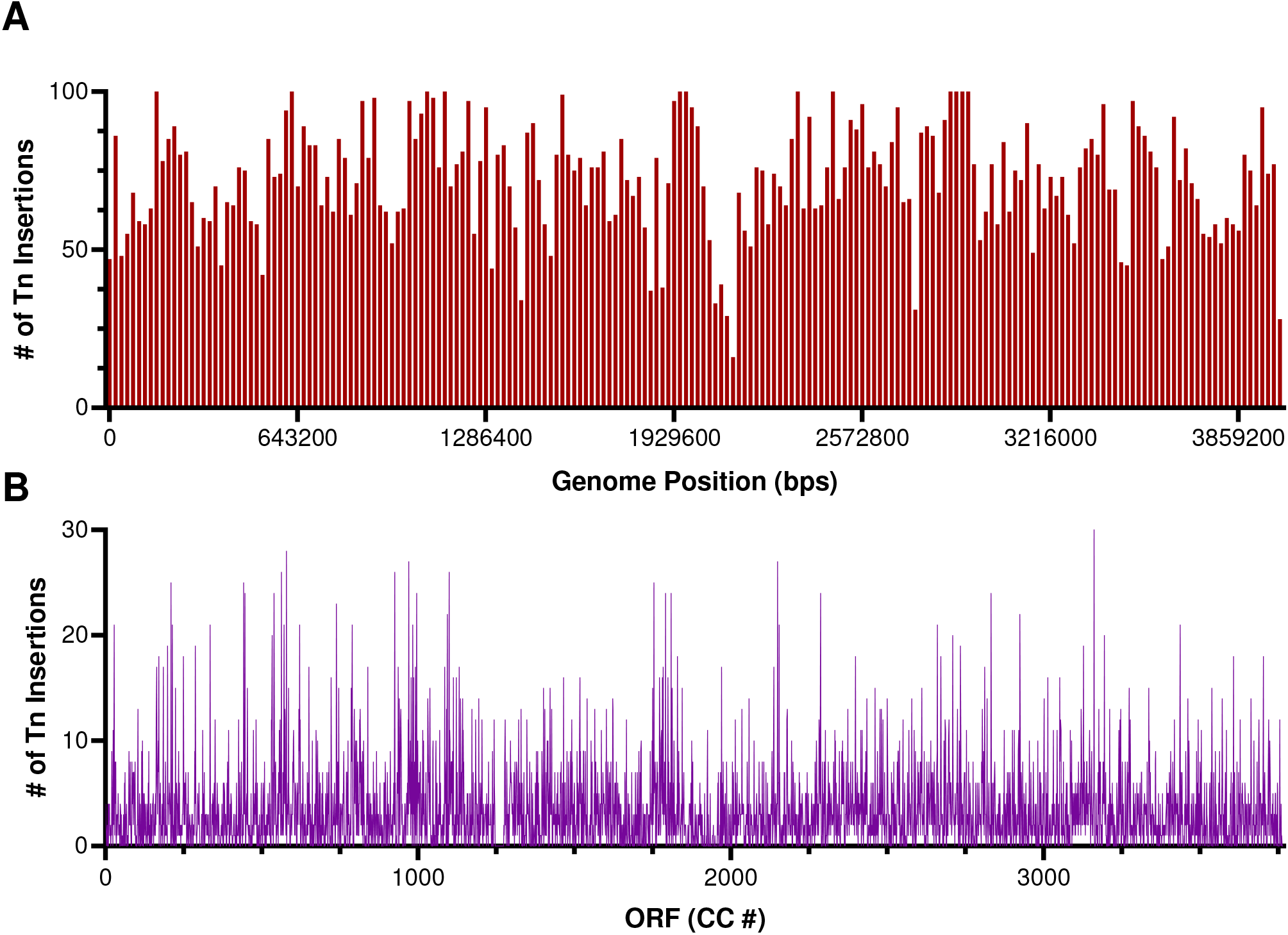
Transposon saturation by A) genome position and B) open reading frame (ORF). Histogram in A) consists of 200 bins with each bin = 20,000 bps.

To generate the CauloKO ordered insertion library, we proceeded to use the Knockout Sudoku approach to determine the plate locations of specific insertion mutants in the initial collection. While the previously-published Knockout Sudoku pipeline was instrumental in constructing the initial collection and performing combinatorial pooling [13], the signal-to-noise in our sequencing data could not be tolerated by the Bayesian inference used in the published Knockout Sudoku software. Thus, we developed our own software to deconvolve the pooled sequencing results. Specifically, we generated a MATLAB GUI that enabled us to semi-automatically determine the predicted coordinates of our mutants based on primary and secondary thresholding requirements of the sequencing read counts (Supplemental Figure 2). The primary threshold was filtered by the absolute number of read counts for a given coordinate. Secondary thresholding determined the signal-to-noise ratio (SNR) of read counts by comparing the highest value for a particular coordinate to the next highest value, as the highest number of read counts was typically the most accurate coordinate of the mutant. Automated filtering by absolute read counts and SNRs provided a high-confidence set of mutants with known locations based on their coordinates. Determining the identity of these mutants with high confidence also enabled us to manually eliminate their positions as possible locations for other mutants, thereby enabling us to triangulate the positions of additional mutants with fewer known coordinates from the initial analysis. The combination of automated and manual mutant selection using the CauloKO GUI enabled us to determine unique genomic insertion sites and plate coordinates for 20,345 mutants, or about ~70% of total mutants picked (Table 1).

**Table 1:** SNPs present in parental strain and transposon mutant from library collection. Rows highlighted in green reflect shared identity between strains. SNPs unique to the parental strain or transposon mutants are highlighted in blue or yellow, respectively.

|  | ORFs Hit | ORFs Missing | Total |
| --- | --- | --- | --- |
| Non-Essential (%) | 2731 (87.1%) | 404 (12.8%) | 3135 |
| High Fitness (%) | 118 (75.6%) | 38 (24.4%) | 156 |
| Essential (%) | 376 (79.0%) | 100 (21.0%) | 476 |
| Total (%) | 3225 (85.6%) | 572 (14.4%) | 3767 |

### CauloKO provides broad coverage of the non-essential genome of C. crescentus CB15

The 20,345 mutants whose positions in the initial collection could be uniquely identified mapped to 2,902 genes that we consolidated into an ordered CauloKO library. CauloKO represents 77% of all predicted ORFs, including 86% of non-essential genes, 18% of essential genes, and 19% of “high fitness” genes (genes previously shown to tolerate transposon mutagenesis at lower rates that other non-essential genes [26]) (Table 1). We hypothesized that the presence of insertions in essential and high fitness genes in our library was due to either strain-specific differences in essentiality between CB15 and NA1000, or retention of partial gene function upon transposon mutagenesis. Retention of partial gene function would be expected for insertions at the ends of ORFs (where enough gene product is retained to maintain cell viability) or the beginnings of ORFS (where alternative start sites or readthrough from transposon promoters could maintain expression of most of the gene). To test this hypothesis, we mapped transposon insertion sites relative to the fraction of the gene into which each transposon inserted. As expected, we found that insertion sites in non-essential genes were uniformly distributed throughout the genes, while insertions in essential genes were biased to the gene ends (Figure 3). Altogether, only 6 genes previously-annotated as essential contained insertions in the middle of the gene. Given this low number and the fact that some of the six genes with central insertions might be essential in NA1000 but not CB15, our results suggest that our library’s coverage and annotation accuracy are robust.

**Figure 3:**
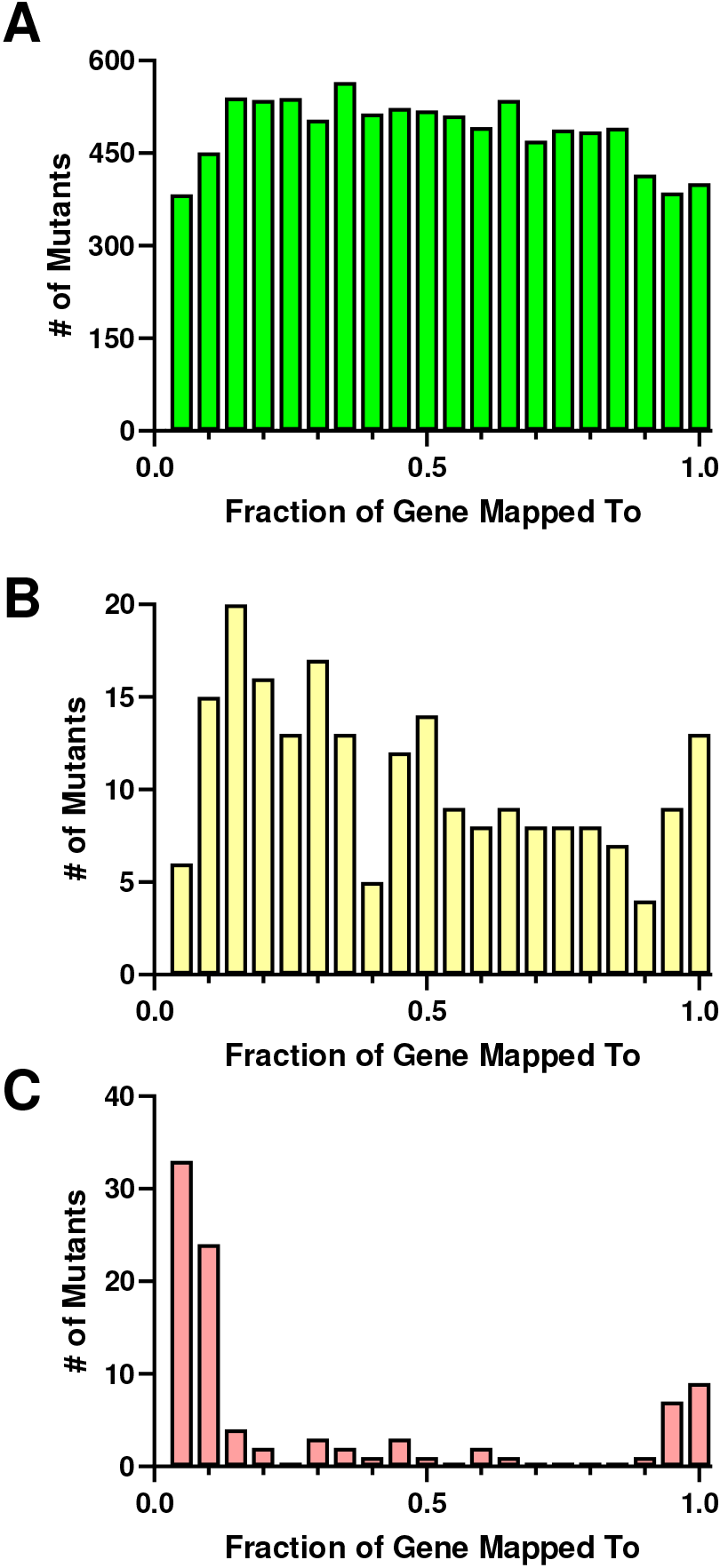
Fraction of gene possessing transposon insertion for essential genes in CauloKO library. A) Nonessential genes (n = 9749). B) High fitness genes (n = 214). C) Essential genes (n = 96). Each bin represents 0.05 fractions of the gene.

### Genetic verification of select mutants and CauloKO portal for community annotation

To confirm the accuracy of our transposon mutant annotations, we used random priming to independently sequence the insertion sites of 53 randomly-selected transposon mutants from the CauloKO library. Our results indicated that 40 of the 53 mutants (75%) contained insertions in either the predicted gene or the gene immediately adjacent (Supplemental Table 2). Past efforts have noted that transposon library creation leads to a fraction of inaccurately annotated mutants and this misannotation rate is not known for many libraries [3–7]. This problem has been addressed by creating redundant library collections when possible: with a 75% accuracy rate, the chance that at least one of two insertions in a given gene is correct is 94% and for genes with three insertions that chance rises to 99%. Thus, we included mutants in triplicate when possible in the final collection, leading to the final CauloKO library of 6739 mutants depicted in Supplemental Table 2. In this final collection, 54% of the ORFs were represented by 3 independent insertions, 25% by 2 insertions, and only 21% by one insertion (Table 2). Together, we expect that 93% of the ORFs in this collection are accurately represented by at least one insertion in the corresponding gene (Supplementary Table 3). We also included several mutants with insertions in intergenic genomic regions as internal controls, as these do not create disruptions to predicted gene products but do contain a transposon. For ease of screening more complex phenotypes, we also created a “consolidated” collection with a single mutant for each ORF, with the representative mutant having a transposon inserted at the most upstream possible position to favor disruption of the gene product (Supplemental Table 4).

**Table 2:**
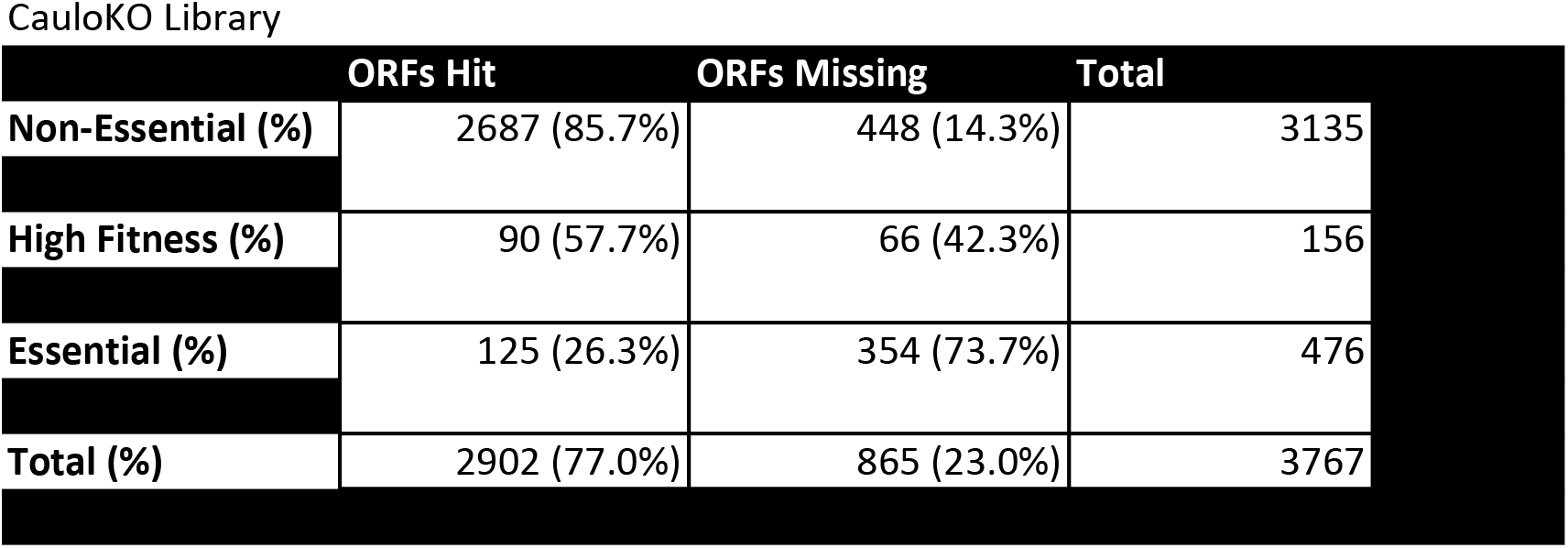
Number of insertion mutants and genetic mutants present in single, duplicate, or triplicate in CauloKO.

CauloKO Library
|  | ORFs Hit | ORFs Missing | Total |
| --- | --- | --- | --- |
| Non-Essential (%) | 2687 (85.7%) | 448 (14.3%) | 3135 |
| High Fitness (%) | 90 (57.7%) | 66 (42.3%) | 156 |
| Essential (%) | 125 (26.3%) | 354 (73.7%) | 476 |
| Total (%) | 2902 (77.0%) | 865 (23.0%) | 3767 |

No library is perfect and future work by the community can help to significantly improve CauloKO and its annotation. Consequently, we developed a mechanism for members of the community that use CauloKO to update the rest of the community in real-time as to the identity and properties of mutants in the collection. This is a problem with other ordered libraries, as annotations are rarely updated when passed to a new user. Specifically, we created an online portal to allow for individuals from any lab to add updated gene annotations for the CauloKO library from Sanger sequencing or other sequencing methods to confirm annotations (cauloko.princeton.edu).

### Phenotypic validation by crystal violet screening

Using the consolidated collection, we wanted to demonstrate the utility of the CauloKO library in a phenotypic screen that would both validate known genes and identify novel genes with a particular phenotype. For this purpose, we assayed our library for biofilm formation using crystal violet staining for quantification [29–31]. While surface attachment has been robustly examined in *C. crescentus* and is known to be involved in biofilm formation [32], crystal violet screening has not been previously performed in *C. crescentus*. Thus, known surface attachment mutants should be identified by our screen as positive controls. Furthermore, biofilm formation is a population-level phenotype such that this assay highlights the power of screening our library in clonal populations.

Surface attachment is mediated by motile swarmer cells that use Type IV Pili (TFP) for initial adhesion. Retraction of the pilus initiates cell cycle regulation, transforming swarmer cells into surface-adherent stalked cells [33–35]. This irreversible attachment is mediated by the holdfast, an incredibly strong adhesin that is synthesized intracellularly, exported to extracellular space, and anchored to the end of the stalk [36–39]. Once attached, microcolony formation and biomass production lead to larger-scale biofilm structures. Holdfast synthesis and anchoring, pilus activity, and flagella presence are required to promote higher order biofilm formation [32]. Additionally, extracellular DNA from *C. crescentus* can inhibit biofilm formation by blocking holdfast attachment to surfaces [40]. While each of these processes has been suggested to be involved in biofilm formation, the lack of previous systematic biofilm screens left unclear the extent to which each of *C. crescentus’* attachment modalities contributes to biofilm development.

To establish the power of CauloKO for genetic screens and phenotypic analysis of mutants of interest, we screened the consolidated CauloKO library for biofilm formation by measuring normalized crystal violet staining after 48 hours of growth in 96-well plates. Plates were stained, measured for crystal violet (by OD_540_), and normalized for bacterial growth (by OD_660_) to determine biofilm presence in biological triplicate. Mutants with repeated decreased or increased biofilm formation relative to the average biofilm formation of their respective plate are listed in Table 3.

**Table 3:** Mutants identified in crystal violet screen of CauloKO library.

| Redudnancy | # of Mutants |
| --- | --- |
| Single | 625 |
| Double | 717 |
| Triple | 1560 |
| Total | 6739 |
| Intergenic | 65 |

We first determined whether the mutants we identified were consistent with previous findings. Chemotaxis mutants are known to stimulate biofilm formation while the stalk development mutant, *podJ*, is known to reduce biofilm formation, and we found that these mutants affected crystal violet staining in the expected direction in our screen (Figure 4B) [41, 42]. Holdfast synthesis was also previously shown to be required for crystal violet staining in exponentially-growing cells. Consistently, most genes annotated as essential for holdfast formation in exponential phase also displayed reduced crystal violet staining in our screen (Figure 4A). Some holdfast-related genes are regulatory and do not eliminate holdfast synthesis itself. In agreement with prior studies, these holdfast-associated mutants that do not reduce the formation of the holdfast itself did not decrease crystal violet staining (Figure 5A) [37, 39]. Across all holdfast-related genes, we found only four whose results from our stationary-phase crystal violet screen did not agree with previous reports from exponential phase studies: glycosyltransferase *hfsG*, holdfast synthesis protein *hfsA*, Polysaccharide polymerase *hfsC*, and holdfast anchor component *hfaC* (Figure 5A) [36–39]. These discrepancies could be due to either incorrect gene annotation or due to biological differences between the specific conditions used for the different assays [32].

**Figure 4:**
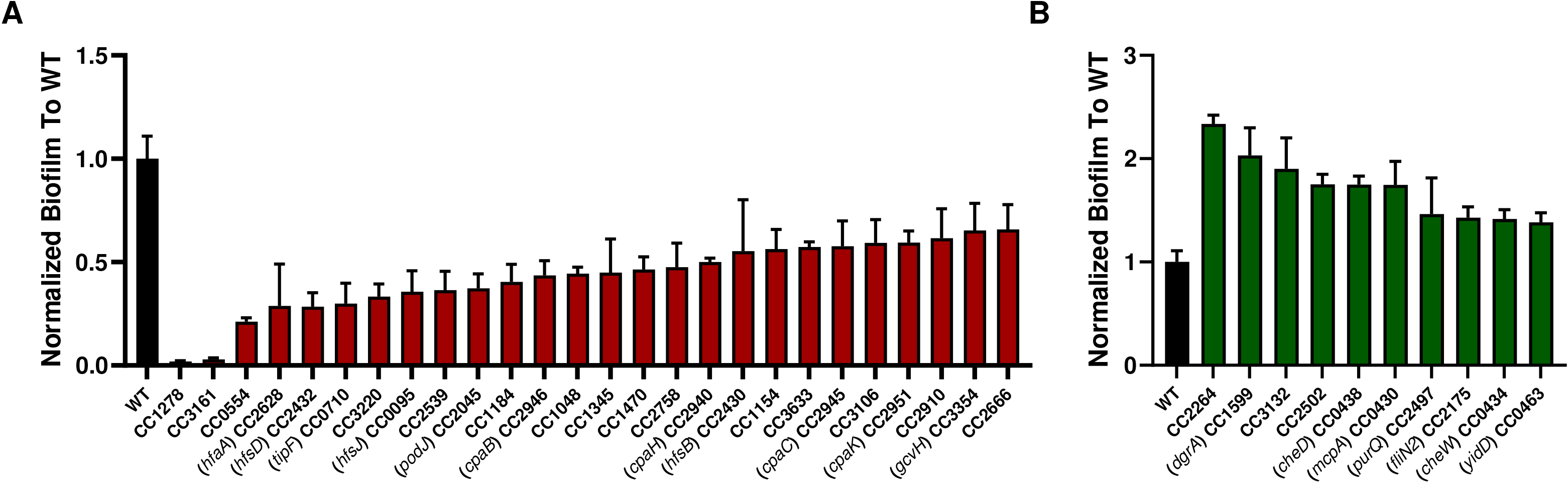
Quantification of significant mutants from crystal violet screen. A) Mutants decreased for biofilm formation by crystal violet staining. B) Mutants increased for biofilm formation by crystal violet staining.

While the role of the holdfast in crystal violet staining is clear, the role of flagella and TFP in biofilm formation is more complicated. Recent work demonstrated that flagellar disruption stimulates holdfast synthesis, which would be predicted to stimulate crystal violet staining in our screen [43, 44]. Consistent with this finding, no flagellar mutants led to decreased biofilm formation in our screen and several flagellar mutants led to increased biofilm formation (Figure 4B, Figure 5C). However, several flagellar mutants retained wild-type levels of biofilm formation, suggesting that there may be specific roles for individual flagellar components in holdfast regulation.

**Figure 5:**
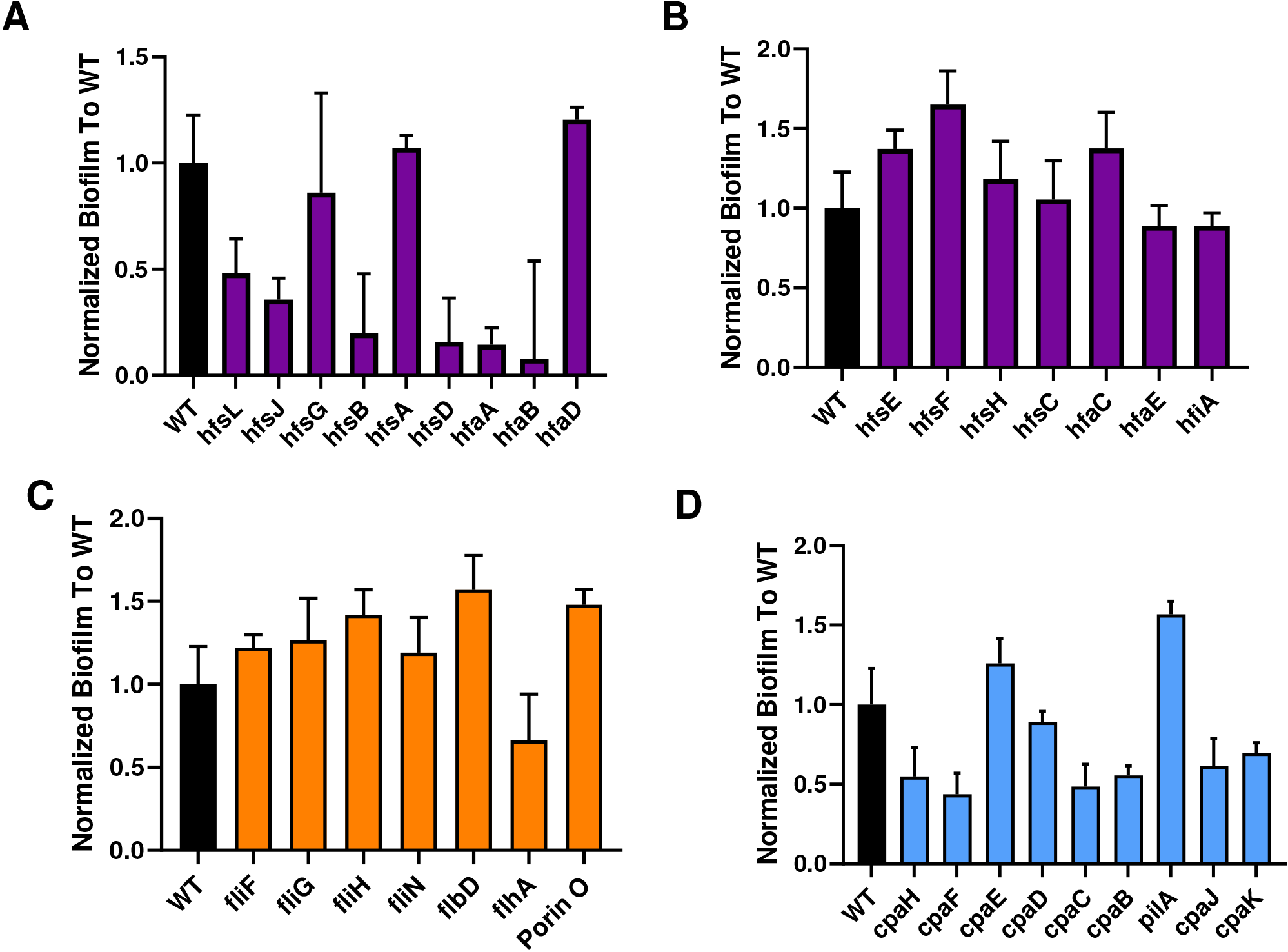
Select mutants related to biofilm formation replicate known phenotypes. Holdfast (A-B, purple), flagella (C, orange), and pilus (D, blue) mutants were measured for crystal violet staining normalized to biomass after forty-eight hours of growth. Error bars represent standard error of the mean. cpaG and cpaA of the Type IV pilus operon not present in CauloKO library.

TFP are extracellular polymers in which PilA pilin monomers are assembled and disassembled by a TFP motor [33]. A recent TnSeq study of *C. crescentus* attachment to cheesecloth found that loss of the *pilA* TFP subunit led to increased cheesecloth attachment while loss of motor components led to decreased cheesecloth attachment, suggesting that unassembled PilA monomers regulate holdfast formation [45]. In our screen, we also saw that *pilA* and TFP motor mutants displayed the opposite phenotypes; we observed a decrease in biofilm formation in *pilA* and an increase in biofilm formation in motor mutants (Figure 4A, Figure 5D). Thus, while further studies will be needed to dissect the regulatory cross-talk and timing of how flagella, TFP, and holdfast structures affect one another and biofilm formation, our results both confirm previously-described regulatory interactions and identify new questions for future work.

Our screening results also identified potential novel players involved in *C. crescentus* biofilm formation (Figure 4). Amongst mutants with decreased crystal violet staining, two mutants stood out as having remarkably low holdfast staining: CC1278, which encodes a predicted GMC family oxidoreductase, and CC3161, which encodes a TonB-dependent receptor. These putative gene functions have not been previously identified as necessary for biofilm formation in *Caulobacter*, but in *Acinetobacter baumannii*, a TonB-dependent copper receptor leads to a dramatic decrease in biofilm formation [46]. Amongst mutants with increased biofilm formation, CC2264, a phosphomannomutase/phosphoglucomutase (PMM/PMG), shows particular promise as a novel biofilm factor. In *Pseudomonas aeruginosa*, AlgC acts as a bifunctional PMM/PMG to regulate exopolysaccharide production and specificity [47]. The only polysaccharide known to be required for biofilm formation in *Caulobacter* is holdfast. It is thus tempting to speculate that PMM/PMG may act on holdfasts or that additional exopolysaccharides may be involved in biofilm formation. CC2264 mutants also displayed increased adherence to cheesecloth [45], supporting the importance of this gene in biofilm development. Additional studies will be needed to determine the specific roles these new candidates play in *Caulobacter* biofilm development.

## Conclusion

Our study presents both a new resource for the community for holistically assessing *C. crescentus* physiology and new data on biofilm formation in this species. Our findings confirm previous results, thereby validating the library and our approach. We also find new mutants and regulatory interactions, raising new mechanistic questions for future studies that one would not have known to address otherwise. The CauloKO library is particularly promising for tackling new genetic screens that require clonal populations or could not otherwise be performed by TnSeq, such as the crystal violet screen presented in this study. Our work also presents a new sequencing analysis approach for decoding barcoded pools of transposon mutants. The CauloKO GUI, though used only for this study at present, allows for visual representation of sequencing data that is user-friendly and more intuitive than other sequencing pipelines and may thus be useful for other applications. We have thus made this GUI open source and available for the community. Finally, we hope that the online CauloKO portal will promote efforts across the *Caulobacter* community to make this library a dynamic and long-lasting resource to enable future discoveries.

## Methods

### Bacterial stains and growth conditions

For this study, an overnight culture is defined as a single colony inoculated in 5 ml tubes and grown for 16 hours. Exponential phase cultures were obtained by a 20-fold back dilution of overnight culture in fresh media and grown to an OD_660_ of ~0.5. *Caulobacter crescentus* laboratory strains (CB15) were grown in shaking culture at 30°C in PYE media and pMiniHimar1 containing *E. coli* were grown as described previously [13, 48].

### Library preparation and sequencing

To generate mutant colonies and sequencing pools for the CauloKO library, we utilized the combinatorial pooling method described in the Knockout Sudoku pipeline [13]. Paired-end 150 nt Illumina MiSeq sequencing was performed on all sequencing pools at Princeton University’s Genomics Core and analyzed using Knockout Sudoku computational tools. For determining the physical location of mutants, we generated a Matlab-based CauloKO GUI for determining sequencing read counts for all mutants in the collection. Functions of the GUI are described in greater detail in Supplementary Figure 2. We used an absolute read count threshold of 50 read counts for primary thresholding and a signal-to-noise ratio (SNR) of 9 for secondary thresholding. Sanger sequencing verification of mutants was accomplished through Genewiz.

### Crystal violet assay

CauloKO mutants were statically grown for 48 hours at 30°C in 96-well plates in ensure saturation of growth. Absorbance (OD_660_) was measured using a plate reader to obtain density of the culture, and then washed with deionized water. Crystal violet (0.5% w/v in 80% dH_2_0, 20% methanol) was added to the plates and incubated at room temperature for 10 minutes before washing again in deionized water. Crystal violet was resolubilized using 95% ethanol as a solvent and absorbance (OD_540_) was a measured using a plate reader. Biofilm production was measured by normalizing the absorbance of the crystal violet to the absorbance of the bacterial growth (OD_540_ / OD_660_). For the crystal violet screen, each consolidated CauloKO library plate was measured in biological triplicate. For targeted candidates in the collection, mutants were measured in biological triplicate with each experiment performed in four technical replicates. The wild-type used for experiments was the *C. crescentus* CB15 parental strain of the library.

## Acknowledgements

We would like to thank the Barstow group at the Department of Chemistry at Cornell University for providing reagents and technical assistance for the CauloKO library. Additionally, we’d like to thank the Princeton University Sequencing Core for providing sequencing services for the CauloKO library. We also wish to particularly thank the many kind lab members and volunteers who assisted in the effort of hand-picking colonies for the CauloKO library: DJ, AW, AS, BG, ES, JC, JW, JM, JS, EA, DV, RS, SC, EB, GV, JS. Finally, we would like to thank our co-authors for taking time to proof-read previous iterations of this manuscript. For researchers interested in utilizing the CauloKO library, please contact the Gitai lab for access.

---

**Supplementary Figure 1: Mutants required for saturation of ORFs as determined by Knockout Sudoku pipeline.** Solid black line represents the total number of ORFs in the *C. crescentus* CB15 genome. Dashed line represents the number of mutants picked for library construction.

**Supplemental Figure 2: CauloKO GUI for sequence data analysis.**

**Supplemental Table 1: Identity of transposon mutants by Sanger Sequencing.** Green indicates matches between Sanger results and library annotation. Yellow indicates partial matching most likely due to arbitrary PCR amplification schema. Red indicates contradiction between Sanger results and library annotation.

**Supplementary Table 2: Identity of transposon mutants by Sanger Sequencing.** Green indicates matches between Sanger results and library annotation. Yellow indicates partial matching most likely due to arbitrary PCR amplification schema. Red indicates contradiction between Sanger results and library annotation

**Supplemental Table 3: Complete CauloKO Library Collection.**

**Supplemental Table 4: Consolidated CauloKO Library Collection.**

| Decreased Biofilm Formation |  | Increased Biofilm Formation |  |
| --- | --- | --- | --- |
| ORF # | Predicted Function | ORF # | Predicted Function |
| 95 | hfsJ | 1184 | Amino acid permease |
| 133 | Ketoacyl reductase hetN | 430 | Chemoreceptor McpA |
| 136 | Uncharacterized protein | 434 | Chemotaxis protein CheW |
| 247 | Two-component receiver protein SpdR | 435 | Chemotaxis protein methyltransferase |
| 451 | Zn-dependent hydrolase, glyoxalase II family | 436 | Protein-glutamate methylesterase/protein-glutamine glutaminase 1 cheB1 |
| 554 | Uncharacterized protein | 438 | Probable chemoreceptor glutamine deamidase CheD |
| 710 | Cyclic-di-GMP phosphodiesterase, flagellum assembly factor TipF | 463 | Putative membrane protein insertion efficiency factor |
| 853 | Uncharacterized protein | 663 | TonB-dependent receptor |
| 898 | Flagellin flgL | 752 | Hypoxia transcriptional regulator FixK |
| 998 | ABC transporter ATP-binding protein | 863 | Uncharacterized protein |
| 1048 | Acylamino-acid-releasing enzyme | 993 | Conserved hypothetical cytosolic protein |
| 1080 | LuxR-family transcriptional regulator | 1599 | Diguanylate receptor protein dgrA |
| 1096 | Ribosomal RNA large subunit methyltransferase J | 1661 | Alanine racemase |
| 1154 | dATP pyrophosphohydrolase | 1700 | Uncharacterized |
| 1184 | Amino acid permease | 1744 | D-alanine aminotransferase |
| 1278 | GMC family oxidoreductase | 2175 | FliN family protein |
| 1345 | TetR/AcrR family transcriptional regulator | 2264 | Phosphomannomutase/phosphoglucomutase |
| 1372 | Uncharacterized protein | 2497 | Phosphoribosylformylglycinamide synthase subunit PurQ |
| 1403 | Cytochrome c oxidase, CcoQ subunit | 2502 | Aminopeptidase |
| 1470 | Acetyltransferase, GNAT family | 2520 | SpoU rRNA methylase family protein |
| 1486 | WbqP Undecaprenyl-phosphate beta-N-acetyl-D-fucosaminephosphotransferase | 2575 | 3-hydroxybutyryl-CoA dehydratase |
| 1641 | 2'-5' RNA ligase family protein | 2604 | Serine-pyruvate aminotransferase |
| 1732 | Ferritin-like domain protein | 3027 | Conserved membrane spanning protein |
| 1737 | Competence/damage-inducible protein CinA | 3083 | NADH-dependent flavin oxidoreductase |
| 1906 | Citrate synthase | 3132 | Acetyl ornithine aminotransferase |
| 2045 | Localization factor PodJL | 3195 | Outer membrane protein oprM |
| 2253 | Lysophospholipase L2 | 3217 | Uncharacterized protein |
| 2385 | Uncharacterized protein | 3237 | Holliday junction ATP-dependent DNA helicase RuvA |
| 2430 | HfsB | 3274 | Multidrug/protein/lipid ABC transporter family, ATP-binding and permease protein |
| 2432 | hfsD | 3346 | Esterase/lipase |
| 2466 | DNA polymerase IV dinB | 3482 | SWIM zinc finger family protein |
| 2539 | M61 glycyl aminopeptidase |  |  |
| 2601 | Cyclopropane fatty acid synthase-like protein |  |  |
| 2628 | hfaA |  |  |
| 2629 | Putative transcription activator protein HfaB |  |  |
| 2642 | Choline dehydrogenase |  |  |
| 2666 | uncharacterized |  |  |
| 2758 | Periplasmic serine endoprotease DegP-like |  |  |
| 2910 | Nucleoside/nucleotide kinase |  |  |
| 2928 | TonB-dependent receptor |  |  |
| 2940 | TadC-related pilus assembly protein CpaH |  |  |
| 2945 | Pilus assembly protein CpaC |  |  |
| 2946 | Pilus assembly protein CpaB |  |  |
| 2951 | TadE-related pilus assembly protein CpaK |  |  |
| 3106 | Alanine racemase |  |  |
| 3217 | Uncharacterized protein |  |  |
| 3220 | Uncharacterized protein |  |  |
| 3266 | RNA polymerase sigma factor, SigJ family |  |  |
| 3380 | 2-deoxy-D-gluconate 3-dehydrogenase |  |  |
| 3412 | Acetyltransferase, GNAT family |  |  |
| 3572 | Carbonic anhydrase |  |  |
| 3633 | dTDP-4-dehydrorhamnose 3,5-epimerase |  |  |
| 3639 | FtsZ-localized protein A (fzIA) |  |  |
| 3705 | Outer-membrane lipoprotein carrier protein |  |  |

## References

1. Shuman HA and Silhavy TJ. The Art and Design of Genetic Screens: Escherichia coli. Nature Reviews Genetics. 2003;4(6): 419–431.

2. Pollock DD and Larkin JC. Estimating the Degree of Saturation in Mutant Screens. Genetics. 2004;168(1): 489–502.

3. Liberati NT, Urbach JM, Miyata S, Lee DG, Drenkard E, Wu G, Villanueva J, Wei T, Ausubel FM. An ordered, nonredundant library of Pseudomonas aeruginosa strain PA14 transposon insertion mutants. PNAS. 2006;103(8): 2833–2838.

4. Cameron DE, Urbach JM, Mekalanos JJ. A defined transposon mutant library and its use in identifying motility genes in Vibrio cholerae. PNAS. 2008;105(25): 8736–8741.

5. Held K, Ramage E, Jacobs M, Gallagher L, Manoil C. Sequence-Verified Two-Allele Transposon Mutant Library for Pseudomonas aeruginosa PAO1. Journal of Bacteriology. 2012;194(23): 6387–6389.

6. Fernandez-Martinez LT, Del Sol R, Evans MC, Fielding S, Herron PR, Chandra G, Dyson PJ. A transposon insertion single-gene knockout library and new ordered cosmid library for the model organism Streptomyces coelicolor A3(2). Antonie Van Leeuwenhoek. 2011;99(3): 515–22.

7. Gallagher LA, Ramage E, Weiss EJ, Radley M, Hayden HS, Held KG, Huse HK, Zurawski DV, Brittnacher MJ, Manoil C. Resources for Genetic and Genomic Analysis of Emerging Pathogen Acinetobacter baumannii. Journal of Bacteriology. 2015;197(12): 2027–35.

8. Baba T, Ara T, Hasegawa M, Takai Y, Okumura Y, Baba M, Datsenko KA, Tomita M, Wanner BL, Mori H. Construction of Escherichia coli K-12 in-frame, single-gene knockout mutants: the Keio collection. Mol Syst Biol. 2006;2: 2006.0008.

9. Cain AM, Barquist L, Goodman AL, Paulson IT, Parkhill J, van Opijnen T. A decade of advances in transposon-insertion sequencing. Nature Reviews Genetics. 2020;21: 526–540.

10. Erlich Y, Chang K, Gordon A, Ronen R, Navon O, Rooks M, Hannon GJ. DNA Sudoku--harnessing high-throughput sequencing for multiplexed specimen analysis. Genome Res. 2009;19(7): 1243–53.

11. Cao CC and Sun X. Combinatorial pooled sequencing: experiment design and decoding. Quantitative Biology. 2016;4: 36–46.

12. Vandewille K, Festjens N, Plets E, Vuylsteke M, Saeys Y, Callewaert N. Characterization of genome-wide ordered sequence-tagged Mycobacterium mutant libraries by Cartesian Pooling-Coordinate Sequencing. Nature Communications. 2015;6: 7106.

13. Anzai IA, Shaket L, Adesina O, Baym M, Barstow B. Rapid curation of gene disruption collections using Knockout Sudoku. Nature Protocols. 2017;12: 2110–2137.

14. Shiver AL, Culver R, Deutschbauer AM, Huang KC. Rapid ordering of barcoded transposon insertion libraries of anaerobic bacteria. Nature Protocols. 2021.

15. Henrici T, Johnson DE. Studies of Freshwater Bacteria. J Bacteriol. 1935;30(1):61–93.

16. Poindexter JS. Biological properties and classification of the Caulobacter group. Bacteriol Rev. 1964;28: 231–95.

17. Ausmees N, Kuhn JR, Jacobs-Wagner C. The bacterial cytoskeleton: an intermediate filament-like function in cell shape. Cell. 2003;115(6): 705–13

18. Skeker JM and Laub MT. Cell-cycle progression and the generation of asymmetry in Caulobacter crescentus. Nature Reviews Microbiology. 2004;2: 325–337.

19. Wilhelm RC. Following the terrestrial tracks of Caulobacter - redefining the ecology of a reputed aquatic oligotroph. The ISME journal. 2018;12(12):3025–37.

20. Moore, G.M. and Gitai, Z. (2020) Both clinical and environmental Caulobacter species are virulent in the Galleria mellonella infection model. PLOS ONE, 15(3): e0230006.

21. West L, Yang D, Stephens C. Use of the Caulobacter crescentus Genome Sequence To Develop a Method for Systematic Genetic Mapping. Journal of Bacteriology. 184(8): 2155–2166.

22. Werner JN, Chen EY, Guberman JM, Zippilli A, Irgon JJ, Gitai Z. Quantitative genome-scale analysis of protein localization in an asymmetric bacterium. PNAS. 2009;106(19): 7858–7863

23. Lasker K, Schrader JM, Men Y, Marshik T, Dill DL, McAdams HH, Shapiro, L. CauloBrowser: A systems biology resource for Caulobacter crescentus. Nucleic Acids Res. 2016;44(D1):640.

24. Marks ME, Castro-Rojas CM, Teiling C, Du L, Kapatral V, Walunas TL, Crosson, S. The genetic basis of laboratory adaptation in Caulobacter crescentus. J Bacteriol. 2010;192(14):3678–88.

25. Kirkpatrick CL and Voilier PH. Synthetic Interaction between the TipN Polarity Factor and an AcrAB-Family Efflux Pump Implicates Cell Polarity in Bacterial Drug Resistance. Chem Biol. 21(5): 657–65.

26. Christen B, Abeliuk E, Collier JM, Kalogeraki VS, Passarelli B, Coller JA, Fero MJ, McAdams HH, Shapiro L. The essential genome of a bacterium. Mol Syst Biol. 2011;7: 528.

27. Nierman WC, Feldblyum TV, Laub MT, Paulsen IT, Nelson KE, Eisen JA, Heidelberg JF, Alley MR, Ohta N, Maddock JR, Potocka I, Nelson WC, Newton A, Stephens C, Phadke ND, Ely B, Deboy RT, Dodson RJ, Durkin AS, Gwinn ML, Haft DH, Kolonay JF, Smit J, Craven MB, Khouri H, Shetty J, Berry K, Utterback T, Tran K, Wolf A, Vamathevan J, Ermolaeva M, White O, Salzberg SL, Venter JC, Shapiro L, Fraser CM. Complete genome sequence of Caulobacter crescentus. Proc Natl Acad Sci USA. 2001;98(7):4136–41.

28. Erhardt H, Steimle S, Muders V, Pohl T, Walter J, Friedrich T. Disruption of individual nuo-genes leads to the formation of partially assembled NADH:ubiquinone oxidoreductase (complex I) in Escherichia coli. Biochimica et Biophysica Acta (BBA). 2012;1817(6): 863–871.

29. Feoktistova M, Geserick P, Leverkus M. Crystal Violet Assay for Determining Viability of Cultured Cells. Cold Spring Harbor Protocols. 2016.

30. Merritt JH, Kadouri DE, O’Toole GA. Growing and Analyzing Static Biofilms. Curr Protoc Microbiol. 2005;0(1): Unit–1B.1.

31. Gilbert-Girard S, Savijoki K, Yli-Kauhaluoma J, Fallarero A. Optimization of a High-Throughput 384-Well Plate-Based Screening Platform with Staphylococcus aureus ATCC 25923 and Pseudomonas aeruginosa ATCC 15442 Biofilms. Int J Mol Sci. 2020;21(9): 3034.

32. Entcheva-Dimitrov P, Spormann AM. Dynamics and control of biofilms of the oligotrophic bacterium Caulobacter crescentus. J Bacteriol. 2004;186(24):8254–66.

33. Ellison CK, Kan J, Dillard RS, Kysela DT, Ducret A, Berne C, Hampton CM, Ke Z, Wright ER, Biais N, Dalia AB, Brun YV. Obstruction of pilus retraction stimulates bacterial surface sensing. Science. 2018;358(6362): 535–538.

34. Medico LD, Cerletti D, Schachle P, Christen M, Christen B. The type IV pilin PilA couples surface attachment and cell-cycle initiation in Caulobacter crescentus. PNAS. 2020;117(17): 9546–9553.

35. Snyder RA, Ellison CK, Severin GB, Whitfield GB, Waters CM, Brun YV. Surface sensing stimulates cellular differentiation in Caulobacter crescentus. PNAS. 2020;117(30): 17984–17991.

36. Smith CS, Hinz A, Bodenmiller D, Larson DE, Brun YV. Identification of genes required for synthesis of the adhesive holdfast in Caulobacter crescentus. Journal of Bacteriology. 2003;185(4): 1432–42.

37. Toh E, Kurtz HD, Brun YV. Characterization of the Caulobacter crescentus holdfast polysaccharide biosynthesis pathway reveals significant redundancy in the initiating glycosyltransferase and polymerase steps. Journal of Bacteriology. 2008;190(21): 7219–31.

38. Berne C, Ellison CK, Agarwal R, Severin GB, Fiebig A, Morton RI, Waters CM, Brun YV. Feedback regulation of Caulobacter crescentus holdfast synthesis by flagellum assembly via the holdfast inhibitor HfiA. Mol Microbiol. 2018;110(2): 219–238.

39. Sulkowski NI, Hardy GG, Brun YV, Bharat, TAM. A Multiprotein Complex Anchors Adhesive Holdfast at the Outer Membrane of Caulobacter crescentus. Journal of Bacteriology. 2019;201(18): e00112–19.

40. Berne C, Kysela DT, Brun YV. A bacterial extracellular DNA inhibits settling of motile progeny cells within a biofilm. Mol Microbiol. 2010;77(4): 815–829.

41. Hershey DM, Fiebig A, Crosson S. A Genome-Wide Analysis of Adhesion in Caulobacter crescentus Identifies New Regulatory and Biosynthetic Components for Holdfast Assembly. mBio. 2019;10(1): e02273–18.

42. Sommer JM and Newton A. Turning Off Flagellum Rotation Requires the Pleiotropic Gene pleD: pleA, pleC, and pleD Define Two Morphogenic Pathways in Caulobacter crescentus. Journal of Bacteriology, 1989;171(1): 392–401.

43. Hershey DM, Fiebig A, Crosson S. Flagellar Perturbations Activate Adhesion through Two Distinct Pathways in Caulobacter crescentus. mBio. 2021;12(1): e03266–20.

44. Berne C and Brun Y. The Two Chemotaxis Clusters in Caulobacter crescentus Play Different Roles in Chemotaxis and Biofilm Regulation. Journal of Bacteriology. 2019;201(18):e00071–19.

45. Hinz AJ, Larson DE, Smith CS, Brun YV. The Caulobacter crescentus polar organelle development protein PodJ is differentially localized and is required for polar targeting of the PleC development regulator. Molecular Microbiology. 2003;47(4): 929–941.

46. Abdollahi S, Rasooli I, Gargari SLM. The role of TonB-dependent copper receptor in virulence of Acinetobacter baumannii. Infect Genet Evol. 2018;60: 181–190.

47. Wei Q and Ma LZ. Biofilm Matrix and Its Regulation in Pseudomonas aeruginosa. Int J Mol Sci. 2013;14(10): 20983–21005.

